# Giant and Opposite Lithium Isotope Effects on Rat Hippocampus Synaptic Activity Revealed by Multi-Electrode Array Electrophysiology

**DOI:** 10.1101/2025.08.23.671929

**Authors:** Khadijeh Esmaeilpour, Irina Bukhteeva, Brian Kendall, Michel J.P. Gingras, Zoya Leonenko, John G. Mielke

## Abstract

Lithium (Li) has been a frontline medication for the treatment of bipolar disorder for decades, but its mechanism of action remains poorly understood^1,2^. While clinically prescribed Li salts consist of a mixture of two stable isotopes, ^6^Li and ^7^Li, the neurobiological effects of each isotope are only beginning to receive attention. Recent theoretical proposals have suggested that the Li isotopes may exert unique effects in the brain, stemming from quantum phenomena linked to their distinct nuclear spin properties^3–5^. However, aside from earlier observations of isotope-dependent variations in animal behavior^6,7^, direct experimental evidence is lacking. We used multi-electrode array electrophysiology to probe field excitatory post-synaptic potentials (fEPSPs) in rat brain slices and demonstrated that ^6^Li and ^7^Li have large and opposite effects on synaptic transmission and lead to smaller, but still distinct, differences in synaptic plasticity, particularly, paired-pulse facilitation and the early stages of long-term potentiation.

Our results provide direct experimental evidence that Li isotopes exert remarkably different neurological effects, while avoiding many of the interpretive challenges associated with behavioural studies. These findings lay the groundwork for future investigations of quantum phenomena in neuronal activity and promote the idea that Li isotopes are pharmacologically distinct agents.

## INTRODUCTION

The dramatic neurological effects of lithium salts were discovered by Cade in the 1940s^8^. Since the mid-1970s, lithium has been a leading medication in the treatment of bipolar disorder, and its potential benefits have been explored in other neurological disorders, including Alzheimer’s disease^1,9,2^. However, in spite of established medical use and considerable research effort over the past seventy years, a complete understanding of the mechanisms of lithium action remains elusive^10,11,1,2^.

Although much research has been pursued to understand the effects and mechanisms of natural lithium salts on the brain and behaviour, clinically prescribed salts consist of two stable nuclear isotopes, ^6^Li and ^7^Li (7.49% and 92.51%, respectively), and only a few studies have explored the neurological effects of the individual lithium isotopes. Of these reports, work done at the cellular and biochemical levels has shown differences that, when observed at all, were small^12–15^. However, animal studies have revealed distinct effects of the lithium isotopes on traits such as locomotor activity and maternal behaviour^6,7,16^, which raises the intriguing question of whether ^6^Li and ^7^Li can affect neuronal function in unique ways.

The possibility that different nuclear isotopes of various elements may play a significant role in biological systems has recently begun attracting interest within the field of quantum biology^17,18^. Of note, two recent theoretical proposals^3,5^ have suggested that ^6^Li and ^7^Li could cause different neurological outcomes due to quantum effects arising from their distinct nuclear spin properties (^7^Li is a spin-3/2 nucleus, whereas ^6^Li is a spin-1 nucleus^19^).

To narrow down the search for lithium isotope effects on the brain, we used acutely prepared rat hippocampal slices, which provide a well-established model for investigating neuronal activity upon exposure to various chemicals and drugs^20^. This approach makes possible adding a well-controlled quantity of lithium while measuring key indicators of neural function (synaptic transmission and synaptic plasticity) in real time to provide experimental results clearly demonstrating a distinct action by each lithium isotope. To our knowledge, such studies have never been pursued before this work. Here, we report a large, highly reproducible, and opposite effect of the two lithium isotopes on both synaptic transmission and synaptic plasticity of rat hippocampal slices measured in real time using multi-electrode array (MEA) electrophysiology. These results indicate that the two lithium isotopes play distinct roles in brain synaptic activity.

## RESULTS

### Lithium isotopes have large and opposite effects on synaptic transmission

Given previous research showing that the acute application of natural Li can alter basal synaptic transmission within rodent-derived brain slices (see Supplementary Material for a review on this matter), we studied the effects of the two stable isotopes of Li on neural activity. To proceed, we measured field excitatory postsynaptic potentials (fEPSPs, or field potentials) evoked from the CA1 region of brain slices acutely prepared from the hippocampus of 8-week old male rats (**Figure 1**) in the presence of the following Li salts: n-LiCl (natural isotopic abundance at 7.49% and 92.51% for ^6^Li and ^7^Li, respectively), ^6^LiCl, ^7^LiCl, n-Li_2_CO_3_, ^6^Li_2_CO_3_, or ^7^Li_2_CO_3_ (for chemical analysis, see Supplementary Table 2).

**Figure 1.**
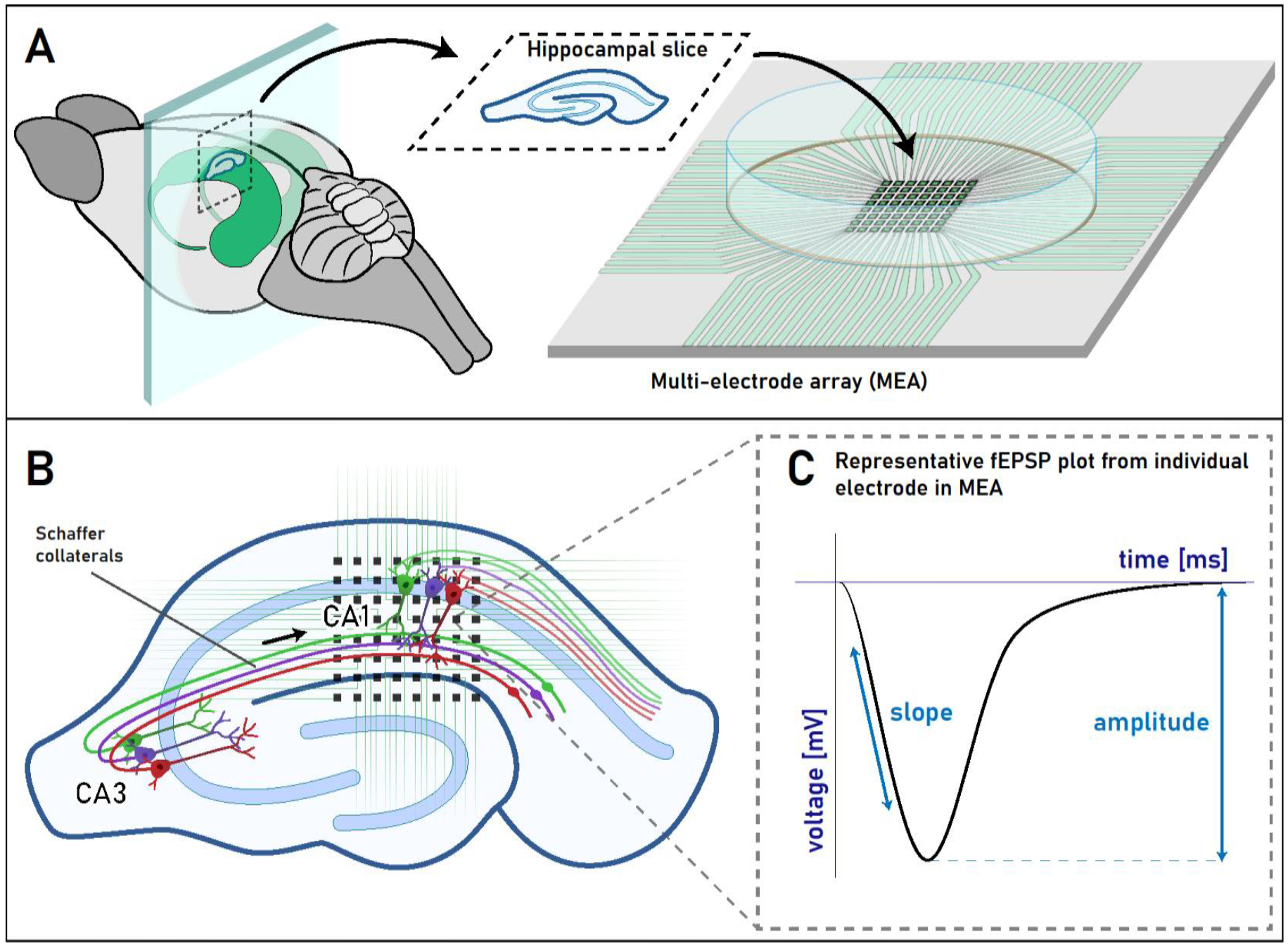
Illustration of the method used to record evoked field potentials within brain slices prepared from the adult rat hippocampus. **(A)** The image displays the relative location of the hippocampus (shown in green) within a typical rat brain. Upon dissection of the hippocampus from the surrounding brain, thin sections (350 µm) were prepared from the dorsal pole and, following a recovery period, were placed over a planar 64-point multi-electrode array (MEA). **(B)** The drawing presents a typical tissue section taken from the dorsal pole of the hippocampus and illustrates 3 representative pyramidal neurons within the CA3 sub-field and their projections (the Schaffer collaterals) that form synapses with the apical dendrites of pyramidal cells found within the CA1 sub-field. The MEA overlay indicates the typical area of the slice from which field potentials were recorded. **(C)** The waveform depicts a characteristic field potential recorded from the CA1 sub-field following stimulation of the Schaffer collaterals. The vertical line denotes the portion of the waveform used to measure amplitude, which is taken to reflect the density of cells residing in the part of the slice surrounding the recording micro-electrode that have responded to stimulation, whilst the slanted line next to the waveform denotes the portion of the downward deflection used to measure slope, which is taken to reflect the speed with which those cells surrounding the micro-electrode have responded to stimulation.

As described in the Methods Section, hippocampal slices were first perfused with artificial cerebrospinal fluid (ACSF) for 20 min (to establish a baseline), followed by perfusion with either 20 mM of a LiCl salt solution, or 10 mM of a Li_2_CO_3_ salt solution for 20 min, and then a 20 min washout period with ACSF. Perfusion with n-LiCl caused a substantial decrease in both the amplitude (↓37%; **Figure 2**) and slope (↓31%; Supplementary Figure 1) of the fEPSP. The ^7^LiCl also caused a decrease in the size of the fEPSP (↓58% amplitude; ↓69% slope), producing a noticeably stronger decrease than observed with n-LiCl. In contrast to both n-LiCl and ^7^LiCl, perfusion with ^6^LiCl resulted in a large *increase* of the fEPSP (↑32% amplitude; ↑29% slope). Notably, the magnitude of difference between isotopes clearly exceeds the threshold for both statistical and practical significance (^7^LiCl vs. ^6^LiCl, end of LiCl perfusion period; amplitude: t(8) = 481.9, p <.0001, d = 305.7; slope: t(8) = 229.9, p <.0001, d = 146.3), and the pattern observed with the chloride salts matches that seen with the carbonate salts (Supplementary Figure 2). In addition, the time needed to reach the maximal amount of change also clearly differed between the isotopes. Whilst the response to ^6^LiCl was very fast, reaching the saturation point in less than 3 min, the responses to n-LiCl and ^7^LiCl were significantly slower and required about 10 min to achieve the maximal degree of difference (**Figure 2B**).

**Figure 2.**
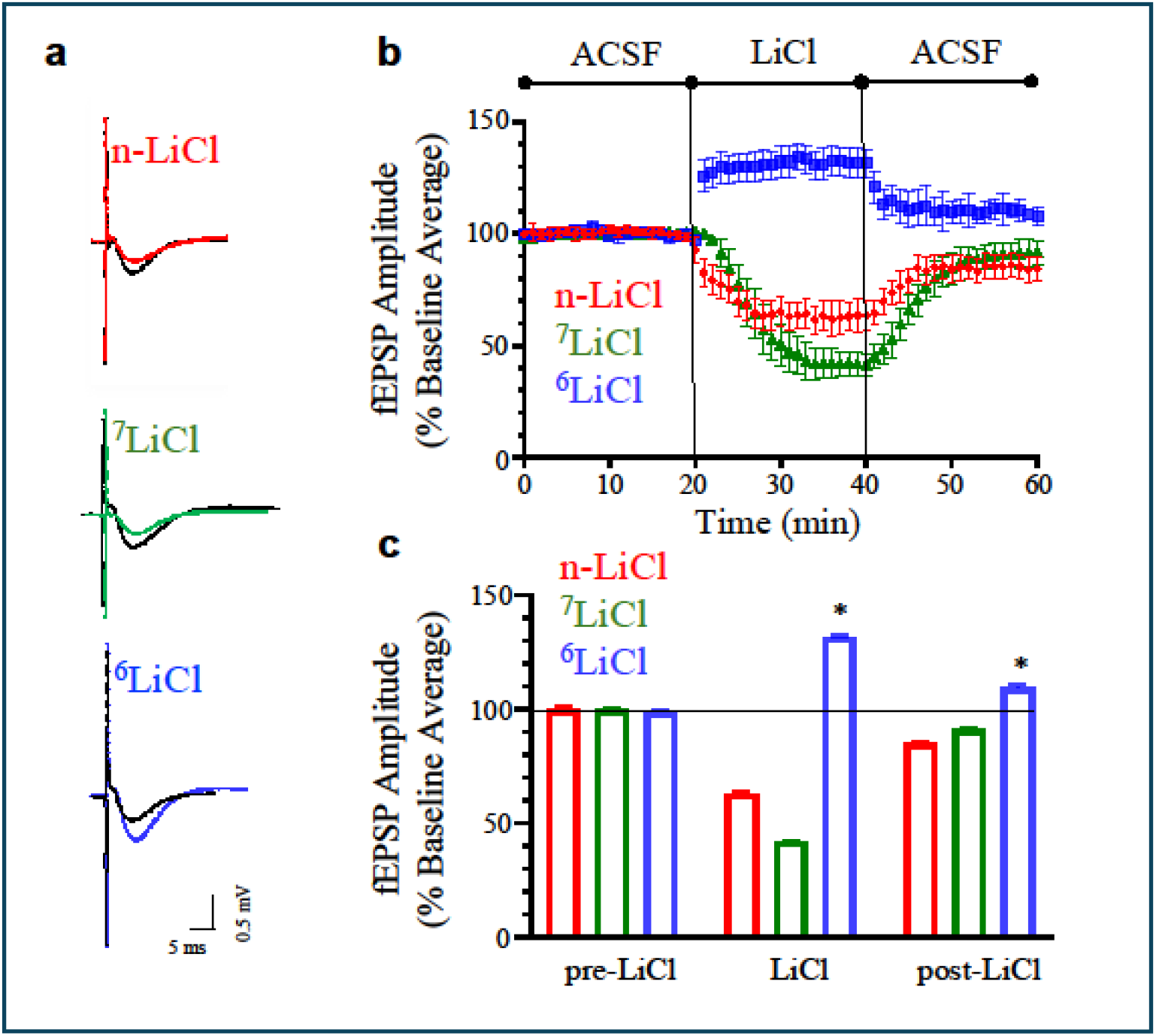
Unique effects of ^6^Li and ^7^Li on basal synaptic transmission. **(A)** Sample fEPSPs recorded from individual slices before (black curves) and after (coloured curves) perfusion with 20 mM of either natural LiCl (n-LiCl), ^7^LiCl, or ^6^LiCl. **(B)** The graph presents the normalised fEPSP amplitudes recorded from slices before and after perfusion with either n-LiCl, or one of the Li isotopes (ACSF, artificial cerebrospinal fluid). Each point shows the mean ± standard deviation (SD) of n = 2 slices from each of N = 5 animals. **(C)** The bar plot illustrates the mean of the normalised fEPSP amplitudes recorded over the last 5 minutes of each experimental phase shown along the x-axis. Each bar in the graph represents the mean ± SD and each asterisk denotes p <.0001 following comparison with the ^7^LiCl group from the same experimental phase.

We thus clearly observed a very large and opposite effect of the two Li isotopes on field potentials, with ^6^LiCl causing a large *increase* and ^7^LiCl a large *decrease* of the fEPSP, with the direction of effect for natural Li salts (approximately 93% ^7^LiCl) resembling that of ^7^LiCl, although the latter caused a stronger decrease. The difference in isotope effects was also observed following washout of the Li salts.

Although none of the responses from any of the groups during the washout period (beginning at the 40 min mark shown in **Figure 2B**) returned fully to their pre-treatment level (the first 20 min of recording), the difference between ^7^LiCl and ^6^LiCl remained clearly visible. At the end of washout, the fEPSP amplitude for slices treated with ^7^LiCl remained lower than baseline by 9%, whilst those treated with ^6^LiCl remained elevated by 10% [t(8) = 30.04, p <.0001, d = 19.05, **Figure 2C**; slope: t(8) = 18.94, p <.0001, d = 11.96, Supplementary Figure 1B]. Importantly, the direction and approximate magnitude of the changes observed were similar regardless of whether LiCl, or Li_2_CO_3_ salts were used (Supplementary Figure 2), which indicates a strong reproducibility in the results.

### Lithium isotopes have significantly different effects on synaptic plasticity

Synaptic plasticity reflects the modifiable nature of connections between nerve cells believed to support the persistent changes needed for memory formation^21,22^ and can be probed with paired-pulse facilitation (PPF), which reflects a short term plasticity on the order of milliseconds^23^, and long-term potentiation (LTP), which reflects plasticity on the order of tens of minutes^22,24^.

#### Paired-Pulse Facilitation (PPF)

We measured PPF in slices before and after perfusion with either n-LiCl, ^7^LiCl, or ^6^LiCl (**Figure 3**). Two stimuli of equal strength were applied to each hippocampal slice 40 ms apart and the percentage increase in the amplitude of the second fEPSP compared to the first fEPSP (**Figure 3A**) was taken as a measure of paired-pulse facilitation [i.e., PPF = fEPSP 2 / fEPSP 1) x 100] both before Li perfusion (PPF 1) and after Li perfusion (PPF 2), as shown in **Figure 3B**.

**Figure 3.**
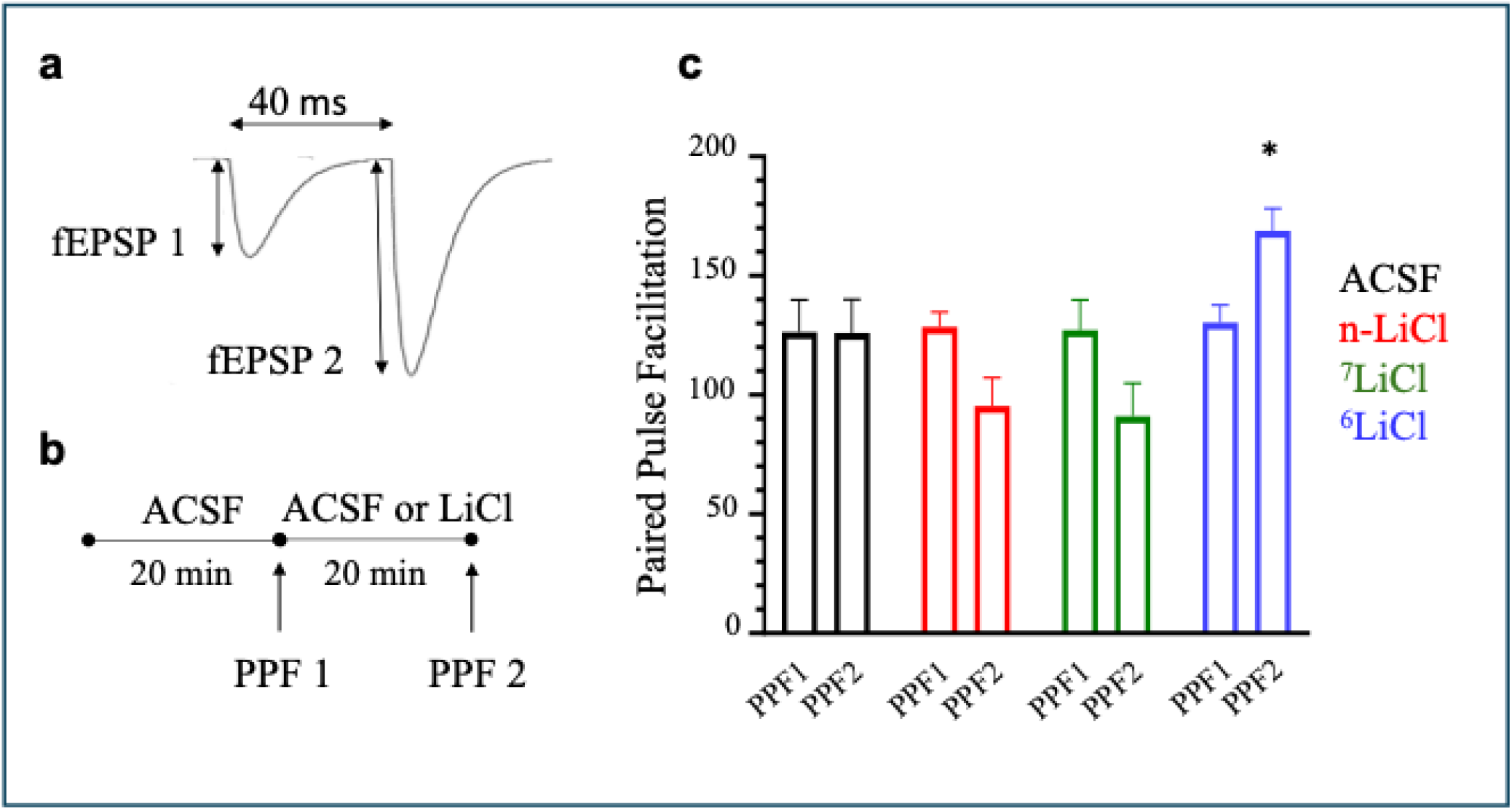
Distinct effects of Li isotopes on paired pulse facilitation (PPF). **(A)** Two stimuli of equal strength were applied 40 ms apart to a hippocampal slice, and the percentage increase in the amplitude of the second fEPSP was taken as a measure of facilitation. **(B)** The experimental timeline illustrates that the magnitude of PPF was measured before and after perfusion with either artificial cerebrospinal fluid (ACSF), 20 mM natural Li, or one of the Li isotopes (20 mM). **(C)** The bar plot shows the magnitude of PPF observed before and after perfusion with either ACSF, or one of the Li-containing solutions. Each bar in the graph represents the mean ± SD of n = 2 slices from each of N = 5 animals; the asterisk indicates p <.0001 relative to PPF 2 from the ^7^LiCl group.

Although the magnitude of PPF observed in both sets of slices was highly similar prior to Li perfusion [PPF 1 signal for the ^7^LiCl group = 127% and PPF 1 for ^6^LiCl group = 130%; t(8) = 0.98, p =.35), a clear difference emerged upon perfusion with the two isotopes. In particular, at the end of the 20 min perfusion period with Li, the PPF 2 for ^7^LiCl was 91% (a clear decrease compared to PPF 1), whilst the PPF 2 for ^6^LiCl was 169% (an obvious increase compared to PPF 1) [**Figure 3C;** t(8) = 14.01, p <.0001, d = 9.39]. Our observations revealed that the application of ^6^LiCl enhanced paired-pulse facilitation, whereas ^7^LiCl depressed this form of plasticity.

#### Long-term potentiation (LTP)

After a 20 min baseline recording period, we perfused slices with Li salts for a further 20 min (which brought about the expected isotope-specific changes described in a previous paragraph), and then used a brief bout of high-frequency stimulation (HFS, or tetanus; a train of 4 pulses at 100 Hz delivered for a duration of 1 s, see Methods) to induce LTP. Following HFS, we measured fEPSPs for an additional 20 min to determine the effect of each isotope on the induction phase of LTP (the first 5 min after HFS, which is also known as post-tetanic potentiation; PTP), or the maintenance phase of LTP (the last 5 min of the recording session, which is also known as short-term potentiation; STP).

Although potentiation was induced in the presence of both isotopes (**Figure 4A**), the magnitude of the change was clearly different between them: a large increase with ^7^LiCl (↑60%) and a much smaller increase with ^6^LiCl: (↑15%) [t(8) = 6.12, p =.0003, d = 3.87; **Figure 4B**]. Whilst we noted a significant difference between the isotopes in their effect upon the induction of LTP, the magnitude of potentiation at the end of the post-HFS recording period (maintenance) was similar for the two isotopes [^7^LiCl: ↑7%; ^6^LiCl: ↑10%; t(8) = 1.61, p =.15; **Figure 4C**].

**Figure 4.**
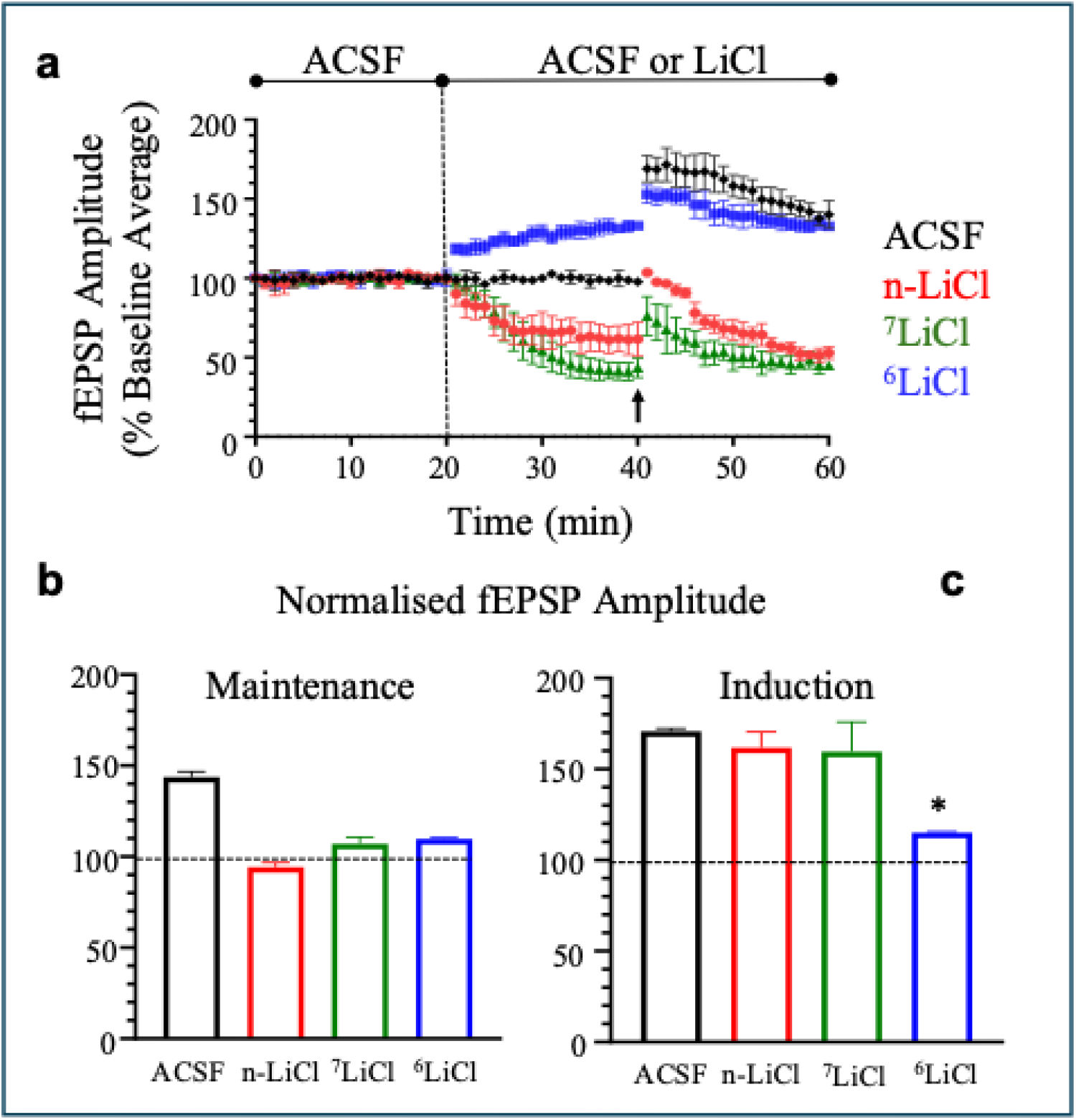
Li isotopes have different effects on long-term potentiation (LTP). **(A)** The graph presents the normalised fEPSP amplitudes recorded from slices perfused with artificial cerebrospinal fluid (ACSF), or a Li-containing solution. Three consecutive periods are shown: baseline (0-20 min), perfusion with ACSF, or a Li-containing solution (20-40 min), and after activation of LTP (40-60 min) by the application of high-frequency stimulation (HFS; denoted by the arrow at 40 min). Each point in the graph shows the mean ± standard deviation (SD) of n = 2 slices from each of N = 5 animals. **(B)** The bar plots illustrate the average degree of change observed in the fEPSP amplitude after application of HFS in either the induction phase (time period from 41 to 45 min), or the maintenance phase (time period from 56 to 60 min) within the different experimental groups. In each case, the changes were measured relative to the final 5 minutes of recording completed before HFS (a period following application of the Li-containing solutions when responses were observed to have become stable). Each bar in the graph presents the mean ± SD of n = 2 slices from each of N = 5 animals; the asterisk indicates p =.0003 relative to ^7^LiCl.

## DISCUSSION

Our results demonstrate that the two Li isotopes have a large and opposite effect on synaptic transmission (**Figure 2**). Specifically, ^6^Li very rapidly increased the strength of synaptic transmission, whereas ^7^Li caused a more gradual, but even greater, decrease in synaptic transmission, similar to n-Li, which is dominated by ^7^Li. The opposite effects of the two Li isotopes on synaptic transmission, together with the rapid reaction time at which a distinct electrophysiological response is observed for ^6^Li, suggest that differences in nuclear spin, rather than mass-dependent factors such as small variations in diffusion constants, underlie the observed phenomena.

Different effects of the Li isotopes were also observed in short-term plasticity –– a large enhancement in PPF by ^6^LiCl and its depression by ^7^LiCl (**Figure 3C**). As well, short-term plasticity induced by high-frequency stimulation revealed distinct patterns, with a large increase seen with ^7^LiCl and a much smaller increase found with ^6^LiCl during the induction phase of LTP (**Figure 4B**, ‘Induction’). Both PPF and the induction phase (*i*.*e*. PTP) are thought to result from rapid pre-synaptic changes in Ca^2+^ levels^25,23^. A much smaller difference was observed between the isotopes in the maintenance phase of LTP (**Figure 4B, ‘**Maintenance’). The changes that occur in the maintenance phase, which is also known as short-term potentiation, have also been suggested to originate in the pre-synaptic terminal^26^, although whether these changes are directly related to Ca^2+^ levels remains to be determined.

Although several previous reports have examined the effects of Li on synaptic activity, all of them used only natural Li (a mixture of ^7^Li and ^6^Li), and the results of these reports cannot be readily compared to our work, or even between each other, due to considerable variability in experimental parameters and settings (Supplementary Material). To provide a clear comparison between the two isotopes, our data demonstrate dramatic differences between ^6^Li and ^7^Li isotopes in synaptic activity measured within the same highly-controlled experimental framework and suggest that each Li isotope may have a unique effect on pre-synaptic function.

There are multiple processes that can be acted upon by Li to affect pre-synaptic activity, including the Na^+^, K^+^ channels involved^10^ in the action potential and ATP-dependent membrane pumps (including Na^+^/K^+^-ATPase^27^), and the Ca^2+^ signaling^28^ machinery, which can influence the release of neurotransmitters from synaptic vesicles. All these pre-synaptic events require energy supplied by mitochondrial bioenergetics^29^.

While effects of n-Li on numerous cellular and molecular targets have been intensely studied^30,31,11,10^, the role of Li isotopes is largely unexplored apart from a few, mostly recent, reports^13–15,32,33^. In particular, Na^+^ channels are known to be affected by n-Li^10^, but show no Li isotope differentiation^33^, while mitochondria revealed noticeable differentiation between Li isotopes with regards to Ca^2+^ sequestration and release^14,15^.

The importance of mitochondria in neuronal activity^34–36^, including synaptic Ca^2+^ signaling and storage, ATP production, production of reactive oxygen species, and the ability to trigger apoptosis^36^, suggests that the large difference we observe in synaptic activities may be a result of specific mitochondrial processes affected by Li isotopes. For example, Li is known to affect mitochondrial function through several other complex pathways^37,38^ including inhibition of glycogen synthase kinase-3beta (GSK-3β)^39^, improving mitochondrial bioenergetics^40,41^ and enhancing the mitochondrial electron transport chain^42^. Lithium ions can also substitute for Mg^2+^ in the ATP-Mg complex^43^, and an effect of Li has been shown on both ATP synthase and the m-PTP channel^44^. However, while mitochondria revealed measurable differentiation in Ca^2+^ uptake and release between Li isotopes^14,15^, it is not via the Li-designated channel Na^+^-Ca^2+^-Li^+^ (NCLX) exchanger^15^, and no isotope differentiation was observed at the level of GSK-3β^13^.

Considering that ^6^Li and ^7^Li differ only in their nuclear mass and spin, Li isotope effects have recently attracted interest within the expanding field of quantum biology^44,17^. With particular relevance to neuronal function, two recent theoretical proposals have suggested that the ^6^Li and ^7^Li isotopes could display different neurological phenomena by affecting Ca^2+^ storage and signaling via inclusion into calcium phosphate clusters, called Posner molecules^3,32^, or through a radical pair (RP) mechanism proposed to be operational in neuronal function^5^. Indeed, Li isotope differentiation was recently observed experimentally in mitochondrial Ca^2+^ cycling^14,15^, as well as in the formation of in vitro Ca phosphate clusters and Posner molecules^32^. Although direct experimental evidence for an RP mechanism affected by ^5^Li is currently lacking, an interesting recent experimental report suggests that such a mechanism may be operating in mitochondrial bioenergetics^45^. While a precise theoretical explanation for our observations is not possible at this time, our data suggest that Li isotopes may either act in a non-trivial and opposing manner upon the same molecular targets, or on different targets, possibly upon mitochondrial processes guiding synaptic activity.

In summary, our results provide the first experimental evidence that Li isotopes have dramatically different effects on synaptic function. These findings may have several important implications as they may help illuminate the pre-synaptic mechanisms responsible for Li’s action as a mood stabiliser and raise new questions within the field of quantum biology about how mass, and/or nuclear spin may be responsible for such effects.

## MATERIALS AND METHODS

### Lithium Salts and Lithium Isotope Purity

Li isotopes and natural Li salts (both LiCl and Li_2_CO_3_, ≥99%) were purchased from Sigma-Aldrich, with their purity further analyzed using in-house inductively coupled plasma mass spectrometry (ICP-MS). All Li salts were dissolved in ultrapure water within a clean laboratory environment and then the concentrations of sodium, magnesium, aluminum, potassium, calcium, titanium, vanadium, chromium, manganese, iron, cobalt, nickel, copper, zinc, arsenic, rubidium, strontium, zirconium, molybdenum, cadmium, antimony, cesium, barium, hafnium, tungsten, rhenium, thallium, lead, thorium, uranium, and mercury in Li isotope samples were quantified using an Agilent 8800 triple quadrupole ICP-MS. For these elements, measured concentrations for ICP and United States Geological Survey water standards are typically within 10% (and always within 25%) of reported values. Impurity concentrations in Li isotope solutions were determined using 10,000 ppb Li solutions. Differences between prepared and experimentally determined values for Li isotope concentrations were analysed using less-concentrated 1000 ppb Li solutions (to avoid exposure of the ICP-MS detector to unnecessarily large Li ion beams). The differences were consistently less than 0.4% across all samples for the more abundant Li isotope in a solution, ensuring the accurate preparation and quantification by ICP-MS. Following ICP-MS analysis, all values were converted to mM to represent the actual concentrations of elements during the experiment (calculated for the case of 20 mM Li addition). Notably, sodium, magnesium, and calcium were not considered contaminants due to their presence in artificial cerebrospinal fluid. The concentration of other contaminants was within the nM range, and their presence, or absence does not correlate with our experimental results. We thus do not attribute any role to contaminants in the observed effects of Li isotopes.

### Experimental Animals

Male, Sprague-Dawley rats (*Rattus norvegicus)* were purchased from Envigo and used at approximately 8 weeks of age (weight 200-250 g). Throughout the study, animals were group-housed in a temperature-controlled room (21°C ± 1°C) on a reverse 12:12 light-dark cycle (lights on at 10:00 p.m.) and allowed free access to both water and standard rodent chow (Teklad 22/5 rodent diet; Envigo). All animals were used in accordance with procedures approved by the University of Waterloo animal care committee, and in agreement with guidelines established by the Canadian Council on Animal Care.

### Hippocampal Slice Preparation

Animals were rendered unconscious by placement in a chamber saturated with CO_2_ (for a period of ∼90 s), and then euthanised by decapitation. Following euthanasia, brains were rapidly removed (∼60 s) and immediately placed in cooled (<4°C) artificial cerebrospinal fluid (ACSF) containing (in mM): 124.0 NaCl (Millipore Sigma, Oakville, ON, Canada; all subsequent reagents from Millipore Sigma, unless otherwise noted), 26.0 NaHCO_3_, 10.0 HEPES, 10.0 glucose, 2.0 CaCl_2_, 3.0 KCl, 1.0 MgSO_4_, and 1.2 NaH_2_PO_4_ and equilibrated with carbogen (95% O_2_/ 5% CO_2_). In all experiments, care was taken to ensure that the pH and osmolality were kept between 7.35-7.45 and 310-320 mOsm, respectively. Next, one hippocampus was extracted (the right hippocampus was preferred; **Figure 1A**) and placed on the platform of a McIlwain tissue chopper (Mickle Laboratory Engineering Co., Surrey, UK); after this, 350 μm thick slices from the dorsal pole were cut and placed upon platforms containing a semi-permeable membrane in separate compartments of an interface incubation chamber (2-3 slices per platform). The ACSF was continuously gassed with carbogen, the incubation chamber was kept at 32.0 ± 0.5°C, and 60-90 minutes were allowed prior to the beginning of experiments.

### Field Potential Recording

#### Basal Synaptic Transmission

Evoked field excitatory postsynaptic potentials (fEPSPs) were recorded by placing a hippocampal slice into a multi-electrode dish (MED; Alpha MED Scientific Inc., Osaka, Japan) such that the CA1 sub-field was located over an 8 x 8 electrode array (electrode size: 50 x 50 µm, inter-electrode distance: 100 µm; **Figure 1B**). Prior to their initial use, MEDs were washed with deionised water and coated overnight with 0.1% polyethyleneimine in 25 mM borate buffer. Plastic wire mesh and a U-shaped weight were gently placed over each slice to improve contact with the microelectrodes. The MED was then connected to a MED64 recording system (Alpha MED Scientific) and continuously perfused with carbogenated ACSF (3 mL/minute, 32°C) while humidified carbogen was directed over the slice.

Following a 20-min stabilisation period, biphasic constant current pulses (0.2 ms) were applied to individual electrodes along the Schaffer collateral pathway to identify the optimal recording site, as defined by the size and shape of the evoked waveforms (**Figure** 1C; typically, stimulation sites and recording electrodes were within 200 µm of one another). Upon selection of a stimulating electrode, an input-output curve (with stimuli generally applied in the range of 5 μA – 40 μA, Supplementary Fig.3) was completed to determine the stimulation intensity required to evoke a response eliciting 30-50% of the maximal fEPSP amplitude (slices in which the maximal amplitude was less than 0.5 mV were not used for experiments). Next, the baseline activity was recorded for a minimum of 20 min (stability of basal responses was monitored and only those slices with less than ∼10% shift were included in subsequent experimental steps). In each recording session, stimulation was applied once every 60 s with the fEPSPs sampled at a frequency of 20 kHz.

Once a stable baseline had been established, either 20 mM of natural LiCl (or one of the stable isotopes: ^7^LiCl, or ^6^LiCl), or 10 mM of natural Li_2_CO_3_ (or one of the stable isotopes: ^7^Li_2_CO_3_, or ^6^Li_2_CO_3_) was perfused for 20 min, which was followed by a further 20-minute washout period using ACSF. The LiCl working solution was made using a 0.5 M stock solution prepared in deionised water, and the Li_2_CO_3_ working solution was made using a 100 mM stock solution prepared in deionised water. Regardless of the Li salt used, neither the osmolality, nor the pH of the working solutions moved beyond our standard ranges for ACSF. To confirm that the act of changing the solutions perfusing a brain slice would not affect synaptic transmission, field recordings were completed before and after perfusion with either 20 mM NaCl, or 10 mM Na2CO3 and were found to be stable (Supplementary Figure 4).

In this work, we used 20 mM Li as the preferred concentration. Earlier work that examined how acutely applied natural Li might affect synaptic transmission in hippocampal slices employed a wide range of concentrations (5-30 mM; see Supplementary Table 1). Preliminary work by our group determined that 5 mM LiCl (either isotope), the lowest concentration used previously, did not alter evoked fEPSPs to an appreciable degree, and that 10 mM altered fEPSPs to only a modest degree (∼15%; data not shown). However, all slices treated with 20 mM LiCl (natural abundance, or either isotope) showed a clear and pronounced change in the evoked fEPSPs that developed over a 20 min period.

#### Synaptic Plasticity

Two forms of synaptic plasticity were examined. The first was paired-pulse facilitation (PPF), which is a form of short-term plasticity that involves an increase in the size of the second of two fEPSPs evoked by stimulations separated by a short inter-stimulus interval (40 ms). Once a stable 20 min baseline had been established for each slice, PPF was measured; after this, either ACSF, or one of the Li-containing solutions was perfused for an additional 20 min and then PPF was measured again. The second form of plasticity examined was long-term potentiation (LTP), which reflects a relatively longer change in the strength of synaptic communication caused by the application of a brief period of high-frequency stimulation (HFS; a train of 4 pulses at 100 Hz delivered for a duration of 1 s). After recording a stable 20 min baseline for each slice, either ACSF, or one of the Li-containing solutions was perfused for an additional 20 min; next, HFS was applied, and then an additional 20 minutes of recording was completed. Regardless of the form of plasticity examined, the stimulation intensity was consistent throughout each slice recording.

### Statistical Analyses

Each experimental group contained at least 5 animals with at least 2 slices used per animal. To assess whether the data in our groups followed a normal distribution and had homogeneous variances, we examined their descriptive statistics (mean, median, and skewness) and graphical summaries (boxplots and Q-Q plots). Based on our assessment, we felt that our data satisfied the primary assumptions required for the use of parametric tests.

The first question we wished to examine concerned whether there was a difference in basal synaptic transmission between the two stable isotopes either during or after their perfusion. As a result, we used a two-tailed, unpaired Student’s t-test to compare the average fEPSP amplitude and slope found with each isotope during the last 5 min of both the perfusion period and the washout period. The second question asked whether the two stable isotopes were able to uniquely affect the magnitude of the PPF observed. As a result, we used a two-tailed, unpaired Student’s t-test to compare the average PPF observed either before, or after a group of slices was perfused with one of the isotopes. Our final question was focused on whether the two stable isotopes differently affected either the induction or maintenance phases of LTP. Consequently, we used a two-tailed, unpaired Student’s t-test to examine the average fEPSP amplitude and slope observed with each isotope during either the first 5 min (induction) or last 5 min (maintenance) of the post-HFS period. Notably, in each case, comparisons were made to the appropriate average fEPSP measurement recorded during the 5 min immediately preceding the application of HFS (by this time in each experiment, changes in the fEPSP following Li perfusion had stabilised). For each comparison, p values were calculated to provide a measure of statistical significance. To assess the practical significance of relevant group differences, we used Cohen’s d values (d), which standardises the difference between group means according to their pooled standard deviation. When interpreting Cohen’s d, we applied the standard conventions for small (d = 0.2), moderate (d = 0.5), and large (d = 0.8) effect sizes.

## ACKNOWLEDGEMENTS

The authors acknowledge funding from New Frontiers Research Fund - Exploration (NFRF-E) program and Quantum Brain Network led by Matthew Fisher (UCSB) and funded by IONIS Pharmaceuticals Inc.; MJPG and BK are supported by the Canada Research Chair program and MJPG and ZL by the University Research Chair program. We acknowledge the help of James Livingstone with sample preparation for testing impurities of Li salts with ICP-MS, and Dr. Brenda Lee and CryoDragon Inc. for preparing Figure 1. We appreciate helpful discussion with Dr. Michael Beazely, Dr. Nicolas Rouleau and all members of the Waterloo Quantum Biology team as well as Dr. Matthew Fisher and the Quantum Brain Network.

