## Supplementary Materials for "Giant and Opposite Lithium Isotope Effects on Rat Hippocampus Synaptic Activity Revealed by Multi-Electrode Array Electrophysiology"

### SUPPLEMENTARY INFORMATION

1. Supplementary Introduction
2. Supplementary Table 1
3. Supplementary Methods
4. Supplementary Table 2
5. Supplementary Figures
6. Supplementary References

### 1. Supplementary Introduction.

#### Structured Summary of Previous Work Examining the Effect of Natural Lithium on Synaptic Transmission in Hippocampal Slices Prepared from Rat Brain

Using a strategy that began with a keyword search through the title and abstract of papers indexed within PubMed and then proceeded with a search through the reference lists found in retrieved articles, we identified nine previous studies that examined the effect of n-Li upon field potentials evoked from hippocampal slices acutely prepared from the rat brain (Supplementary Table 1). Despite sharing the same general research goal, considerable technical variation exists both within these earlier reports and between them and our work. Indeed, the variation in experimental design may be placed into three separate categories: pharmacological characteristics (Li concentration and length of exposure), recording characteristics (hippocampal sub-field and cellular stratum), and animal characteristics (rat strain, animal sex, and animal age). As a result of experimental variation, directly comparing across studies is not possible; however, to permit some degree of evaluation, a focus can be placed on those parameters from each of the three main categories of difference where there is some degree of consistent overlap with our work: Li concentration, hippocampal sub-field, and animal age. When considering reports where the Li concentration was at least half of that used in our work and where the recording site was the same, four studies can be isolated[1–4]; notably, these studies found a rise in field excitatory post-synaptic potential (fEPSP) response following n-Li (range: 26% - 50%), which conflicts with our observation. However, an important technical distinction between these four reports and ours concerns the final parameter, animal age. That is, the reports used animals that were between 2 and 8 weeks of age (although age was not stated by Higashitani et al.[2], the reported weight of the animals was matched with growth curve data for male, Wistar rats to determine an approximate age range of 6-7 weeks[5]).

Although establishing age ranges within the rat lifespan that directly correspond to those in the human is difficult, given numerous inter-species differences, there is consensus that laboratory rats are weaned at about 3 weeks of age, enter adolescence at about 6 weeks of age, and may be regarded as young adults by about 10 weeks of age[6,7]. As a result, those previous reports most methodologically like our study used animals that were in the peri-adolescent range, whereas we used animals that were in young adulthood. Considering that brain regions such as the hippocampus experience changes in synaptic organization over the period extending from peri-adolescence to adulthood, the variability in age across studies may help to explain why our results differ from those of previous reports. For example, subunits of AMPA receptors (a glutamate receptor sub-type that mediates fast excitatory synaptic transmission) display unique changes between post-natal day (PND) 18 and PND 50; that is, whereas the number of GluA1-stained cells decreases by ~30%, the number of GluA2-stained cells rises by ~25%[8]. In addition, specific subunits of NMDA receptors (another glutamate receptor sub-type that also contributes to fast excitatory transmission) also display distinct developmental profiles; in particular, whereas GluN2A displays a dramatic rise in its expression over the peri-adolescent period, the GluN2B subunit shows an equally clear decline in its expression during the same period[9,10]. As well, PSD-95 (a critical scaffolding protein within synapses containing AMPA and NMDA receptors) has been observed to rise by approximately 40% between PND 35 and PND 180[11].

Taken together, the changes observed in critical components of excitatory synapses during the juvenile period can be reasonably expected to influence neuronal physiology. Indeed, the first few months of the rat lifespan involve changes in (at least) two forms of synaptic plasticity. For instance, long-term depression (a persistent reduction in the size of evoked responses) was observed in hippocampal slices taken from animals during the first 3 weeks of life, but not in those that were older than 5 weeks of age[12]. Similarly, depotentiation (a persistent reduction in a previously potentiated response) was found in the hippocampus of adult animals, but not those that were about 2 weeks of age[13]. Although the precise locus at which n-Li acts upon the synapse remains unclear, since the changes in neuronal transmission it can evoke are influenced by synaptic elements that change during the peri-adolescent period, the age of animals used in this sort of experiment appears to be an important variable.

### 2. Supplementary Table 1.

| Author, Year | Rat Strain | Sex | Age, or Body Weight | Recording Site | natural Li | Length of Exposure | Sample Size | Observed Change |
|---|---|---|---|---|---|---|---|---|
| Colino *et al.*, 1998 | Wistar | both sexes were used; results were not stratified by sex | 2 - 4 weeks | CA1 stratum radiatum | 12 mM | 20 min | 6 slices; number of slices per animal not reported | ~50% ↑ fEPSP slope;<br>~14 minutes required to reach maximal change |
| Evans *et al.*, 1990 | Sprague-Dawley | male | 175 - 350 g | CA1 stratum radiatum | 5 mM | 15-25 min | 5 slices; number of slices per animal not reported | ~65% ↑ fEPSP amplitude;<br>time required to reach maximal change not reported |
| Haas *et al.*, 1982 | Wistar | male | 100 - 150 g | CA1 stratum radiatum | 2 mM | 20-30 min | 7 slices; number of slices per animal not reported | ~10% ↑ fEPSP slope;<br>time required to reach maximal change not reported |
| Higashitani *et al.*, 1990 | Wistar | male | 150 - 300 g | CA1 stratum radiatum and stratum pyramidale | 2, 5, and 10 mM | 10 min | 12 slices (for 5 mM LiCl); number of slices per animal not reported | **stratum radiatum responses:** ~31% ↑ fEPSP amplitude;<br>~5 minutes required to reach maximal change<br>**stratum pyramidale responses:**<br>~8% ↑ population spike amplitude |
| Lacaille *et al.*, 1992 | Sprague-Dawley | male | 150 - 225 g | CA1 stratum pyramidale | 1, 2.5, 5, 10, 20, and 30 mM | 20 min | 3-4 slices per concentration; number of slices per animal not reported | **30 mM LiCl:**<br>~21% ↑ population spike amplitude;<br>~5 minutes required to reach maximal change<br>**20 mM LiCl:**<br>~8% ↓ population spike amplitude;<br>time required to reach maximal change not reported |
| Lim *et al.*, 2005 | Sprague-Dawley | male | 6-7 weeks | CA1 stratum radiatum | 1 mM | 20 min | 6 slices; number of slices per animal not reported | no change in fEPSP slope |
| MacVicar *et al.*, 1981 | Wistar | female | 40 - 70 days | dentate gyrus molecular layer | 25 mM | 20 min | 7 slices; number of slices per animal not reported | ~50% ↓ fEPSP amplitude;<br>~10 min required to reach maximal change |
| Rinaldi *et al.*, 1986 | Sprague-Dawley | male | 4 - 8 weeks | CA1 stratum radiatum and stratum pyramidale | 20 mM | 13-31 min | stratum radiatum responses:<br>1 slice from each of 7 animals<br>stratum pyramidale responses:<br>1-2 slices from each of 4 animals | **stratum radiatum responses:** ~26% ↑ fEPSP amplitude<br>**stratum pyramidale responses:**<br>~27% ↑ population spike amplitude<br><br>time required to reach maximal change not reported for either measurement |
| Valentin *et al.*, 1997 | Wistar | sex not identified | 14 - 30 days | CA1 stratum radiatum | 2, 6, and 18 mM | 10 min | number of slices and animals for each concentration not reported | **2 mM LiCl:**<br>~4% ↑ fEPSP slope<br>**6 mM LiCl:**<br>~22% ↑ fEPSP slope<br>**18 mM LiCl:**<br>~32% ↑ fEPSP slope<br><br>~8 min required to reach maximal change across each concentration |

### 3. Supplementary Methods.

#### ICP-MS Studies to Elucidate the Purity of Li Salts Used in the Present Study

To address potential concerns about contamination within the Li isotope salts used in the current work, we used inductively coupled plasma mass spectrometry (ICP-MS), which is known for its multi-element determination capability, wide dynamic range, and high sensitivity. Sample preparation involved dissolving appropriate amounts of Li salts in ultrapure water within a clean lab environment.

In this study, concentrations of Sodium (Na), Magnesium (Mg), Aluminum (Al), Potassium (K), Calcium (Ca), Titanium (Ti), Vanadium (V), Chromium (Cr), Manganese (Mn), Iron (Fe), Cobalt (Co), Nickel (Ni), Copper (Cu), Zinc (Zn), Arsenic (As), Rubidium (Rb), Strontium (Sr), Zirconium (Zr), Molybdenum (Mo), Cadmium (Cd), Antimony (Sb), Cesium (Cs), Barium (Ba), Hafnium (Hf), Tungsten (W), Rhenium (Re), Thallium (Tl), Lead (Pb), Thorium (Th), Uranium (U), and Mercury (Hg) in Li isotope samples were quantified using Agilent 8800 triple quadrupole ICP-MS. Impurity concentrations in Li isotope solutions were determined using 10,000 ppb solutions. Differences between prepared and experimentally determined values (Supplementary Table 3) for Li isotope concentrations were analysed using less-concentrated 1000 ppb Li solutions (to avoid exposure of the ICP-MS detector to unnecessarily large Li ion beams). The differences were consistently less than 0.4% across all samples for the more abundant Li isotope in a solution, ensuring the accurate preparation and quantification by ICP-MS.

Following ICP-MS analysis, all values were converted to mM to represent the actual concentration of elements during the experiment (calculated for the case of 20 mM Li addition). Notably, Na, Mg, and Ca were not considered contaminants due to their presence in the buffer in which the Li salts were dissolved (that is, artificial cerebrospinal fluid; ACSF). The concentration of other contaminants was within the nM range, and their presence, or absence does not correlate with our experimental results. We do not attribute any role to contaminants in the observed effects of Li isotopes.

To reinforce our findings, we cross-referenced this ICP-MS data with analyses of other Li isotope batches (Supplementary Table 2). The matching contaminant numbers ruled out the presence of toxic contaminants in any specific batch, providing further validation of our results.

**4. Supplementary Table 2.**

**Li salt purity assessment with ICP-MS**

| | $^{7}$LiCl | $^{7}Li_2CO_3$ | $^{6}$LiCl | $^{6}Li_2CO_3$ | n-LiCl | n-$Li_2CO_3$ |
|---|---|---|---|---|---|---|
| **Na** | 7.40E-03 | 4.56E-03 | 2.64E-03 | 4.51E-03 | 1.28E-03 | |
| **Mg** | 5.29E-04 | 8.80E-04 | 4.85E-04 | | | |
| **Al** | | 1.60E-03 | | | | |
| **Ca** | | | | 1.78E-02 | | 1.89E-02 |
| **Ti** | | | | | | 9.26E-04 |
| **V** | 1.01E-04 | | 1.07E-04 | | 1.05E-04 | 1.15E-05 |
| **Cr** | | | | 9.59E-05 | | 2.55E-04 |
| **Mn** | 4.68E-05 | | 3.55E-05 | | 2.84E-05 | |
| **Cu** | 3.99E-05 | | | 4.92E-05 | | |
| **Zn** | 3.74E-04 | | | | | |
| **Rb** | 2.12E-05 | | | | | |
| **Sr** | 2.77E-04 | 2.69E-05 | | | | |
| **Mo** | 2.14E-05 | | | | | 4.68E-05 |
| **Sb** | | 2.34E-05 | | | | 1.49E-05 |
| **Ba** | 7.60E-05 | 5.01E-03 | 3.71E-05 | 1.20E-04 | | |
| **W** | | | | 4.04E-04 | | 2.26E-03 |
| **Pb** | 5.83E-06 | | | | 1.69E-05 | |

Supplementary Table 2. Li salt purity assessment with ICP-MS with concentration of elements presented in $\mu$M.

| | $^{7}$LiCl | $^{7}Li_2CO_3$ | $^{6}$LiCl | $^{6}Li_2CO_3$ | n-LiCl | n-$Li_2CO_3$ |
|---|---|---|---|---|---|---|
| Total Li, ppb | 927.4 | 943.7 | 1012.9 | 1081.7 | 945.3 | 952.2 |
| $^{6}$Li, % | 0.1 | 0.1 | 95.6 | 95.7 | 7.7 | 7.8 |
| $^{7}$Li, % | 99.9 | 99.9 | 4.4 | 4.3 | 92.3 | 92.2 |

Supplementary Table 3. Li salts composition assessment with ICP-MS, Li concentration presented in ppb and Li isotopes presented in %.

### 5. Supplementary Figures

#### Supplementary Figure 1

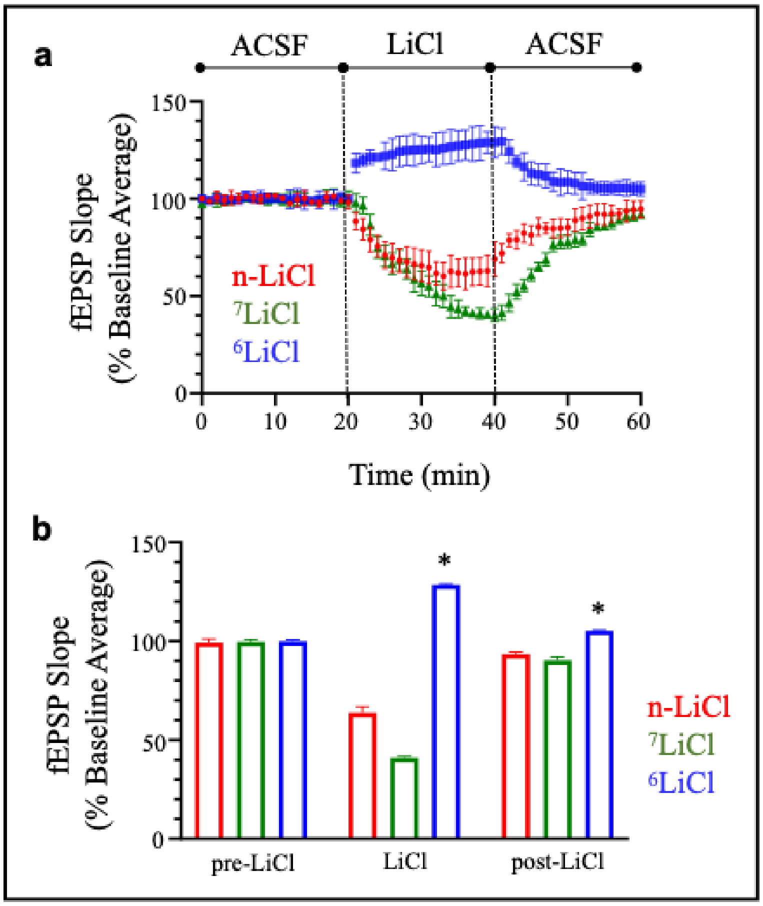


**Supplementary Figure 1.** Unique effects of LiCl and associated lithium isotopes on basal synaptic transmission as measured by fEPSP slope. **(A)** The graph represents the normalised fEPSP slope recorded from slices before and after perfusion with either natural lithium, or one of the lithium isotopes (ACSF, artificial cerebrospinal fluid). Each point shows the mean ± standard deviation (SD) of n = 2 slices from each of N = 5 animals. **(B)** The bar plot illustrates the mean of the normalised fEPSP slopes recorded over the last 5 minutes of each experimental phase shown along the x-axis. Each bar in the graph presents the mean ± SD and each asterisk denotes $p < .05$ following comparison with the $^{7}$LiCl group from the same experimental phase.

**Supplementary Figure 2**

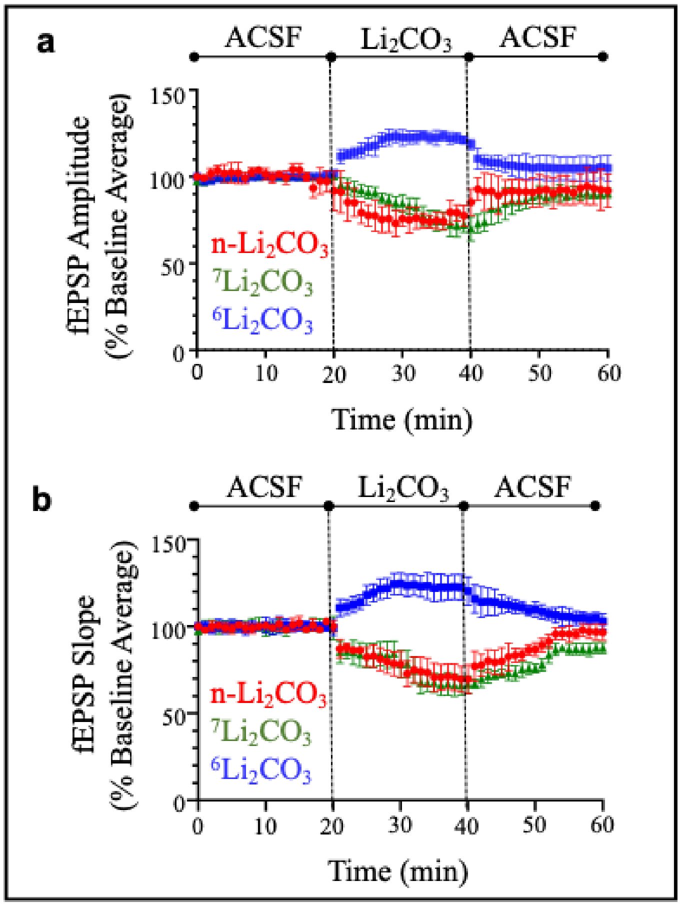


**Supplementary Figure 2**. Differential effects of $Li_2CO_3$ and associated stable lithium isotopes on basal synaptic transmission. The graphs present normalised fEPSP amplitude **(A)** and slope **(B)** recorded from slices before and after perfusion with either $Li_2CO_3$, or one of the stable lithium isotopes (ACSF, artificial cerebrospinal fluid). Each point shows the mean ± standard deviation (SD) of n = 2 slices from each of N = 5 animals.

**Supplementary Figure 3**

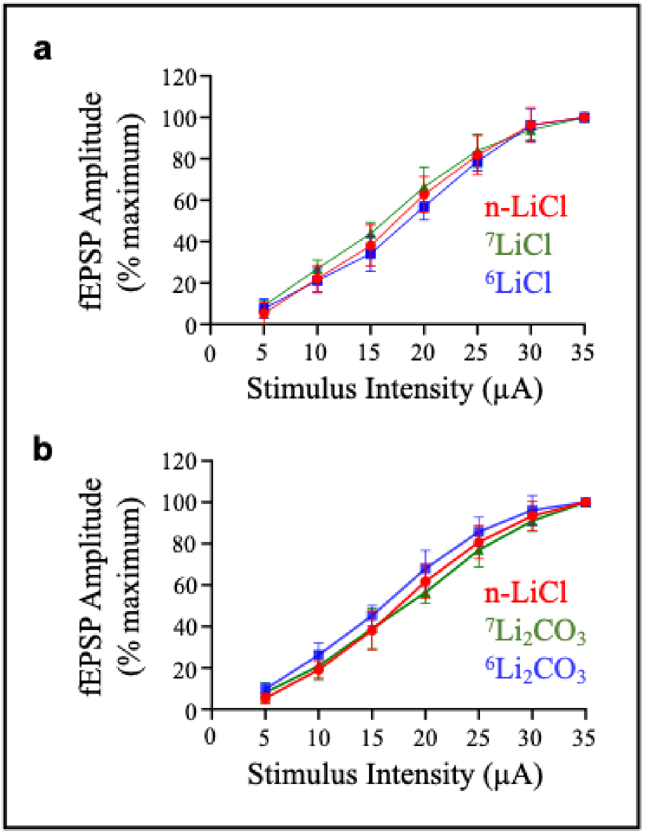


**Supplementary Figure 3**. Synaptic transmission within hippocampal slices appeared similar prior to perfusion with any of the lithium salts. The graphs present stimulus-response curves recorded before perfusion with either 20 mM LiCl **(A)**, or 10 mM $Li_2CO_3$ **(B)**. In each graph, the responses (fEPSP amplitudes) were recorded following incremental increases in stimulation intensity and were normalised to the maximal response recorded for a particular slice (to account for slice-to-slice variation in response magnitude). Each point shows the mean ± standard deviation (SD) of n = 2 slices from each of N = 5 animals.

**Supplementary Figure 4**

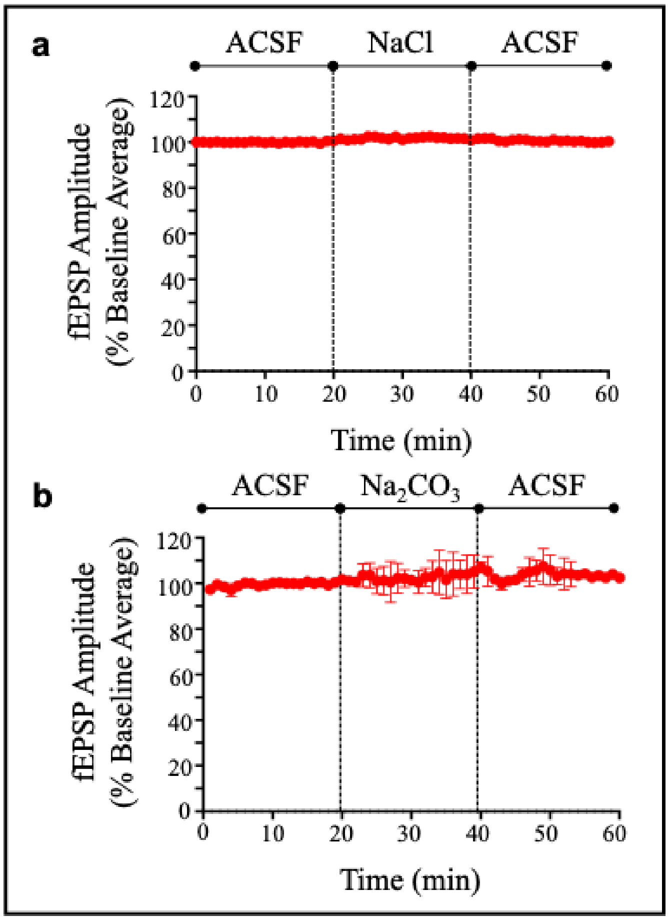


**Supplementary Figure 4**. Substituting either NaCl, or $Na_2CO_3$ in place of natural lithium salts does not affect basal synaptic transmission. The graphs present normalised fEPSP amplitudes recorded from slices before and after perfusion with either 20 mM LiCl **(A)**, or 10 mM $Na_2CO_3$ **(B)** (ACSF, artificial cerebrospinal fluid). Each point shows the mean ± standard deviation (SD) of n = 2 slices from each of N = 2 animals.
